# Host-pathogen co-existence incurs reproductive costs

**DOI:** 10.1101/2020.10.12.336370

**Authors:** Diederik Strubbe, Roel Haesendonck, Elin Verbrugghe, Luc Lens, Richard Ducatelle, Siska Croubels, Freddy Haesebrouck, An Martel, Frank Pasmans

**Author notes:** equal contribution (Diederik Strubbe, Roel Haesendonck). **Abbreviations** CFU: Colony Forming Units; IgY: Immunoglobulin Y; OD: Optical Density; PT: Phage Type.

## Abstract

Widespread endemism of host-adapted pathogens poses a heavy burden on animal and human health. Mechanisms underpinning long-term host pathogen co-existence and concurrent costs are poorly understood. We use infections in pigeons with pathogenic, pigeon adapted *Salmonella* Typhimurium to explain how host and pathogen trade-offs and benefits sustain long-term pathogen endemism. An experimentally infected group of pigeons that was studied for 15 months showed that pathogen persistence decreased host condition and reproductive success, but conferred protection against *Salmonella*-induced clinical disease. The relevance of these findings was confirmed in nature, where this pathogen was shown to widely occur in feral pigeons (*Columba livia*), yet without clinical disease. Pathogen transmission and long-term persistence were associated with intermittent faecal shedding, which markedly increased during crop feeding and natural stress periods. Exploiting host specific traits in the presence of protective host population immunity thus facilitates long-term co-existence, be it at a significant reproductive cost.

## Introduction

While limiting the impact of exposure to pathogens presents a continuous challenge to life on earth, pathogens face a similar struggle to assure their survival and persistence. Globally, bacterial pathogens that depend on a narrow host range as primary niche pose a heavy burden on animal and human health. Pronounced host adaptation requires maximizing pathogen persistence and transmission opportunities while limiting the negative impact on host populations [1,2]. Although key in epidemiology, the mechanisms sustaining endemism of host-adapted bacterial pathogens in the host population and associated trade-offs are currently poorly understood [3,6,8,9] hampering attempts for endemic disease mitigation [6,10]. The underpinning infection dynamics are supposedly driven by host factors (e.g. age, stress and immunity), environment (e.g. biotic and abiotic factors such as food availability, stress, population size and density and pathogen reservoirs) and pathogen determinants such as level of host adaptation, virulence factors and transmission mechanism [3,4]. Adverse effects of endemic infections on the host may vary at the individual and population level [5–7] but overall could be expected to be tempered, avoiding long-term host population declines with substantial reduction of the pathogen’s niche.

We here use *Salmonella* as a model organism to study mechanisms of pathogen endemism. *Salmonella enterica* subspecies *enterica* consists of over 2500 serovars, several of which are known to be endemic in various animal populations and even humans [1,8,11–13]. The host-adapted *Salmonella enterica* subspecies *enterica* serovar Typhimurium (*Salmonella* Typhimurium) PT99 strain in pigeons (*Columba livia*) is characterized by a very narrow host spectrum and pigeons are considered its main host [1,14,15]. Similar to other host-adapted *Salmonella* serovars, the course of an infection with this lineage may vary from subclinical to a typhoid fever-like disease, with a tropism for the host gonads, suggesting that vertical transmission occurs [14,15].

In a one year infection trial, we identified factors driving *Salmonella* infection dynamics during endemism and associated costs and benefits for the host population. We determined to what extent *Salmonella* exploits host specific traits associated with stress (molting, breeding) or reproduction (crop feeding of nestlings, vertical transmission through infected eggs) that may increase bacterial shedding and promote transmission. Finally, we conducted a field study across Flanders (northern Belgium) in which we estimated *Salmonella* prevalence and clinical impact in populations of feral pigeons to confirm the validity of the experimental results.

## Results

### Subclinical *Salmonella* infections are endemic in feral pigeon populations

*Salmonella* Typhimurium occurrence was found in all four studied urban pigeon populations. High risk of exposure to *Salmonella* was demonstrated by marked seroprevalence (33.83%, range: 13.33% – 56.41%, see Fig. 1; average true prevalence varies between 5.0% and 78.4%, 95% confidence interval, see table S1a).

**Fig 1.**
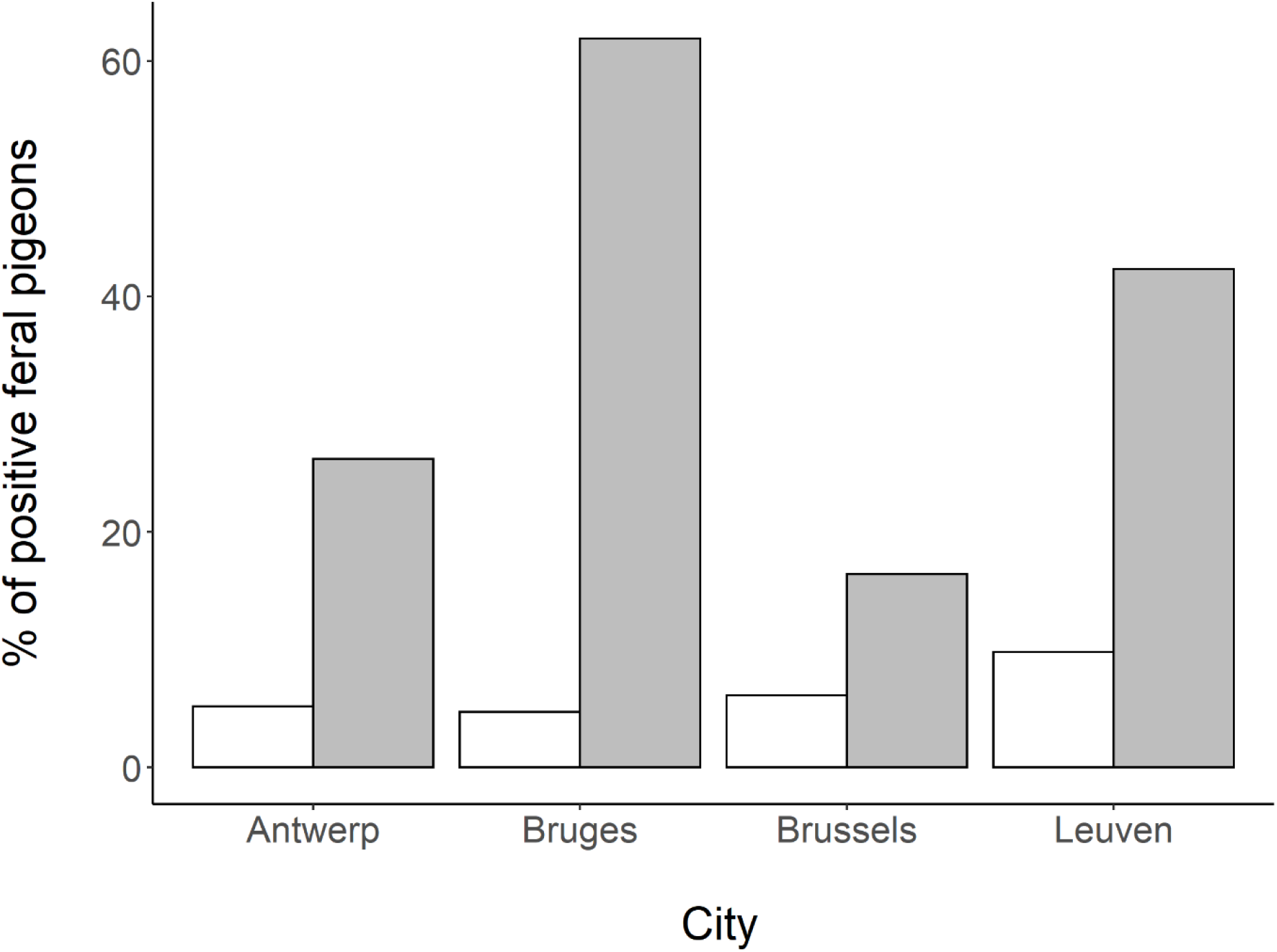
Faecal *Salmonella* shedding and presence of serum *anti-Salmonella* antibodies in feral pigeons. White bars represent the percentage of pigeons from which *Salmonella* Typhimurium was isolated (on average 3.76%), and grey bars the percentage of *Salmonella* seropositive pigeons (on average 33.83%) in a given population.

In all populations, a low percentage of pigeons shed low numbers of *Salmonella* Typhimurium in the faeces (average apparent prevalence: 3.76%, range: 2.56% – 6.90%; average true prevalence varies between 0.8% and 8.4%, 95% confidence interval, table S1b). There was no significant correlation between health status, body constitution score and the presence of faecal *Salmonella* or serum anti-*Salmonella* antibodies within hosts (all P-values > 0.164; see online appendix tables S2a and S2b).

### Disease occurs during the initial phase of infection only, but long-term persistence incurs a reproductive

*Salmonella* Typhimurium infection dynamics in an experimentally infected group of pigeons were followed up during 66 weeks, spanning two consecutive breeding seasons (Fig. 2) and including natural stress periods (molt, breeding, introduction of new group members). Shortly after inoculation, *Salmonella* shedding increased drastically (Fig. 2).

**Fig 2.**
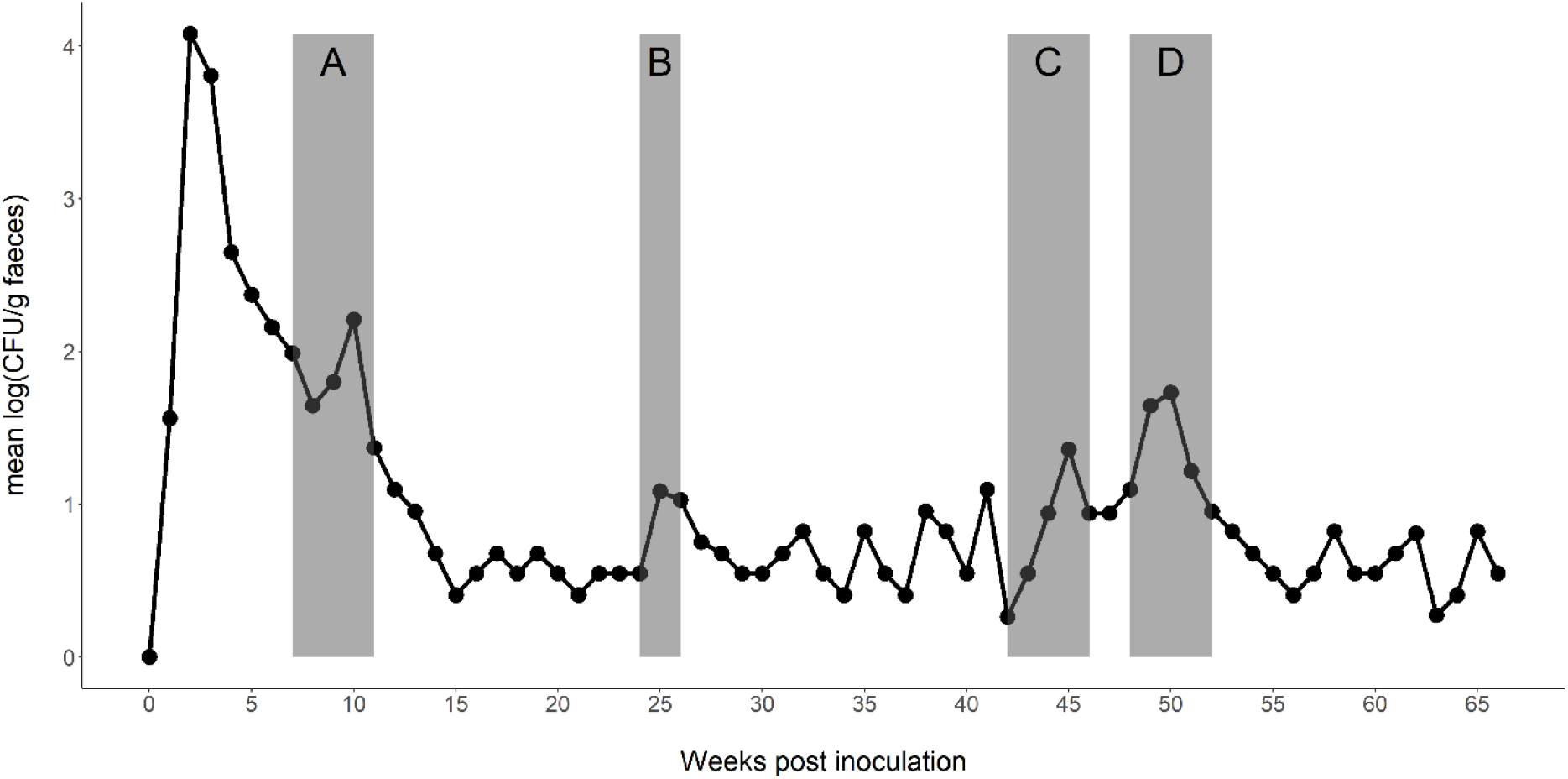
Temporal dynamics of faecal *Salmonella* shedding by experimentally infected pigeons. Experimentally inoculated pigeons showed a significant increase in shedding during: A, breeding period 1 (Parental); B, molting; C, introduction of naïve pigeons and D, breeding period 2 (Parental + F1 generation). Grey bars indicate the duration of each stressful period.

The best fitting model (i.e. the lowest AIC-value: 17.38; table S3) obtained for describing trends in *Salmonella* shedding after the inoculation period had an ARIMA(1,0,1) structure with non-zero mean, and included both induced stress periods and the number of pigeons present as important external regressors (P-values < 0.0001). This first-order autoregressive model (AR1 estimate and standard error: −0.47±0.26) with simple exponential smoothing confirms that after inoculation, *Salmonella* faecal shedding declines towards a nonzero mean (2.27±0.29, P-value < 0.0001; i.e. *Salmonella* remains present in the population). Alleged stress periods correspond to temporary increases in *Salmonella* shedding (0.42±0.08, P-value < 0.0001) but with a negative correlation between the number of pigeons in an aviary and *Salmonella* faecal shedding (−0.04±0.01, P-value < 0.0001; table S3).

*Salmonella* was shed intermittently at low levels in the faeces during the entire experimental trial of 66 weeks (Fig. 2). This continued presence of *Salmonella* and temporary host-stress associated *Salmonella* flare-ups did not result in persistent disease in the chronically infected group during the 66-week experiment. Only shortly after initial and experimental inoculation of the naive founder pigeons, clinical signs of paratyphoid (e.g. diarrhea, anorexia, polydipsia) were observed. Body mass trends differed between the experimentally infected and the control group (interaction between month and treatment, P-value < 0.0001), but this difference is mainly due to lower body masses of experimentally inoculated pigeons in the time periods immediately following the inoculation (experimental infection versus control group month 1: −39.10±7.65, P-value < 0.001; month 2: −23.65±7.70, P-value = 0.012). In later months, body weights of infected birds remained lower than those of control birds (P-values > 0.05 but < 0.10) but only for the last month, this difference was significant (P-value = 0.03, Fig. 3; table S4). Clinical recovery was established 14 days post exposure and no clinical signs of paratyphoid were noted thereafter.

**Fig 3.**
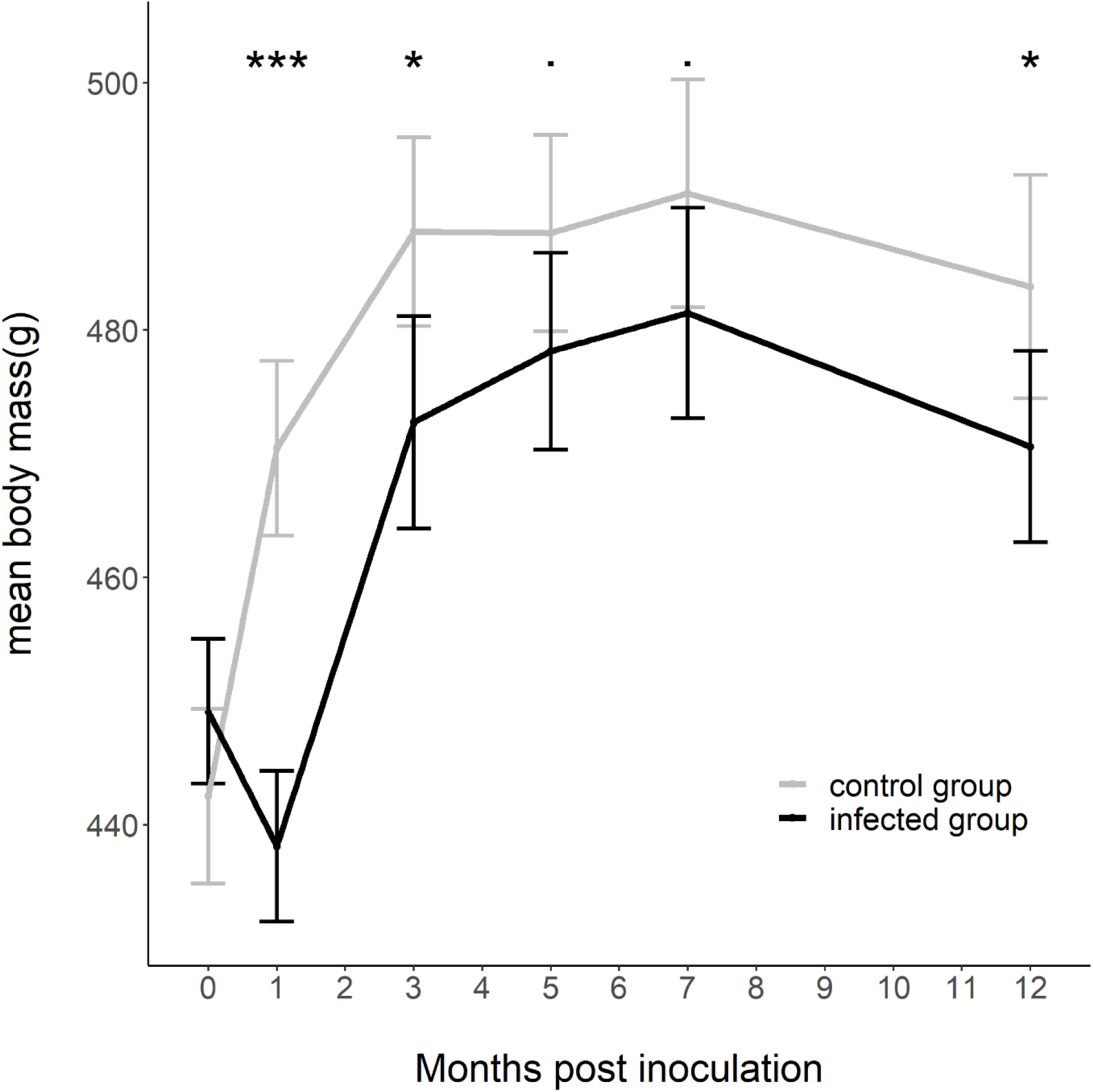
Mean body weight of the adult pigeons in the *Salmonella* infected vs. the negative control group. The mean body weight per group is given (mean ± se) before and after inoculation with *Salmonella* Typhimurium varietas Copenhagen DAB69 and sham inoculation respectively. Significance codes: *** < 0.001, ** <0.01, * < 0.05, 0.05 < . < 0.10.

### *Salmonella* exploits host-specific behavior for horizontal transmission

The crops of six out of eight pigeons that crop-fed their offspring were positive for *Salmonella* after enrichment. All crop-fed nestlings with at least one positive parent became positive for *Salmonella* from day three after hatching. Adult pigeons excreting *Salmonella* in the crop did not simultaneously shed detectable numbers of *Salmonella* in the faeces. During the first nesting period (28 days), no *Salmonella* was detected in any of the cloacal swabs, but during the second breeding period, *Salmonella* was detected in the cloacal swabs of 53.3% (8/15) of the nestlings, with first *Salmonella* detections on average from day 4.6 ± 3.6 post hatching.

### Association with the host reproductive tract does not typically result in vertical pathogen transmission but impairs host reproductive success

*Salmonella* Typhimurium showed a marked association with the pigeons’ gonads, as evidenced by the isolation of *Salmonella* from 5 of the 13 ovaries and/or oviducts and 6 of the 20 testes. Presence of *Salmonella* in the pigeons’ reproductive tissues was confirmed by immunohistochemical staining (Fig. S1a-b). Despite this association with the pigeons’ gonads, *Salmonella* could not be isolated from any of the semen samples (n = 121) collected during the whole experimental trial. *Salmonella* was found in 4.5 % of the eggs laid (i.e. five out of 111) in the infected group. Four of these 5 eggs were found to be non-fertilized. At day of hatch, none of 35 faecal and cloacal swabs sampled from hatchlings in the infected pigeon group were positive for *Salmonella* during both breeding periods. We thus conclude that, despite strong association with the female reproductive tract, vertical pathogen transmission to the offspring is a rare event.

Association of *Salmonella* with the host reproductive tract correlated with reduced reproductive performance through negative effects on both eggs and nestlings. *Salmonella* infected pigeons had nests characterized by lower egg masses, more shape/shell anomalies and a lower hatching success compared to control group pigeons. Hatchlings also had lower body masses, showed delayed fledging and gained less mass in the 28 days after fledging, -compared to the negative control group (all P-values < 0.028, table 1; tables S5a-f). Clutch size, the number of nests with one egg only and the prevalence of infertile eggs did not differ between groups (all P-values > 0.219, table 1; tables S5g-i). During the first breeding period, none of the 20 nestlings died due to paratyphoid in the infected group. During the second breeding period in this group, five out of 15 nestlings died of paratyphoid during the first 28 days after hatching (on average at day 16 ± 9). No clinical signs were observed among the remaining nestlings, except for 1 nestling during the second breeding period which presented depression and right elbow joint arthritis. Breeding phenology was similar too, as no differences were found in the onset of nesting, nest building and mating behavior, nor in laying dates (P-value = 0.889, table 1; table S5j).

**Table 1.** Reproductive parameters measured during breeding periods. All results are given as total number or average number ± SD.

| Parameter | Negative control group | Infected group | p-value |
| --- | --- | --- | --- |
| Clutch size | $9.44 \pm 0.53$ | $8.54 \pm 2.3$ | 0.537 |
| Shape/shell anomalies | 0 out of 86 eggs | 12 out of 111 eggs | **0.011** |
| Number of 1 egg nests | 10 out of 28 nests | 20 out of 36 nests | 0.219 |
| Egg mass (g) | $19.36 \pm 2.21$ | $18.11 \pm 2.28$ | **0.0009** |
| *Salmonella* presence | 0 out of 86 eggs | 5 out of 111 eggs | n.a |
| Infertile eggs | 15 out of 71 eggs | 26 out of 92 eggs | 0.299 |
| Laying date | $7.25 \pm 0.45$ | $7.28 \pm 0.57$ | 0.889 |
| Hatching success | 88.63% | 75.9% | **0.015** |
| Hatchling mass (g) | 17.27 ± 3.67 | 15.41 ± 2.19 | **0.015** |
| Mass gain over 28-day period (g) | 378 ± 46 | 328 ± 61 | **0.005** |
| Nestling mortality (up to day 28) | 96.4% | 74.3% | **0.028** |
| Fledging age (days) | 28.04 ± 0.51 | 30.03 ± 1.28 | **<0.0001** |

### Acquired population immunity protects against clinical disease but not infection

The host immune response to *Salmonella* infection was evidenced by a pronounced humoral response in the adult pigeons: Enzyme Linked ImmunoSorbent Assay (ELISA) results revealed seroconversion of the experimentally infected pigeons from four weeks after inoculation, whereas the control group remained negative. Infected pigeons seroconverted and remained seropositive until the end of the 66-week experiment (P-value = 0.0010; Fig. S2; Table S6a). The offspring of these pigeons was characterized by significantly higher levels of maternal anti-*Salmonella* antibodies (IgY) in their blood at birth as compared to the control group (P-value = 0.0002). While still significantly different, these antibody titers decreased over a 14-day period (day 0 versus day 14: P-value = 0.009; day 14 versus day 28: P-value = 0.177; Tables S6b-c; Fig S3). After an experimental oral challenge with *Salmonella,* age-matched (5-6 months old) pigeons from the infected group showed a significantly better faecal consistency (P-value=0.0001, Table S6d), shed lower amounts of faecal *Salmonella* (P = 0.0257; Table S6e) and had less clinical signs as compared to the pigeons of the control group (P=0.057, Table S6f). Also for the other parameters tested, pigeons born in the infected group tended to be less strongly affected than those from the control group, but these differences failed to reach statistical significance. After the experimental oral challenge with *Salmonella*, pigeons from the infected group did not have more severe organ lesions (−0.083±0.116, P-value=0.486; Table S6g) compared to the pigeons from the control group. The latter tended to seroconvert slower than the juveniles born in the infected pigeon group (−0.670±0.519, P-value = 0.208; Table S6h, Fig S4). Pigeons from the infected group had similar *Salmonella* organ loads (−0.244±0.808, P =0.767; table S6i) as pigeons from the control group. Pigeons from the infected group were characterized by higher body masses compared to control group pigeons (53.5±22.3 g, P=0.031, table S6j), but both groups exhibited a similar body mass trend (decline in mass through time, −1.844±7.148. P=0.798; table S6k).

### Stress periods result in flare-ups of infection

The two breeding periods, moult and the introduction of five naive animals in the group coincided with marked flare-ups of the *Salmonella* infection, as evidenced by significantly increased faecal *Salmonella* shedding (P-value < 0.0001, see ARIMA model above, Fig. 2; Table S3). This higher faecal *Salmonella* excretion was, however, not accompanied by the occurrence of clinical symptoms, but resulted in an efficient transmission of *Salmonella* to newly-introduced pigeons which started to excrete viable bacteria within on average 8 days after introduction in the positive group, though without showing any signs of illness.

To establish a causal relationship between stress and flare-up of infection, pigeons were housed individually. Individual housing resulted in significantly increased faecal *Salmonella* shedding (P-value < 0.0001; Fig 4A, table S7a), and increased the variance in corticosterone levels in the individually housed pigeons compared to the control group (control group: no change in variance, F-test P-value = 0.584, experimental group: increase in variance, F-test P-value < 0.001, table S7b). Mean plasma corticosterone levels increased as well in individually housed pigeons, although this increase failed to reach statistical significance compared to corticosterone trends in the negative group (P-value=0.325, table S7c, Fig. 4B).

**Fig 4.**
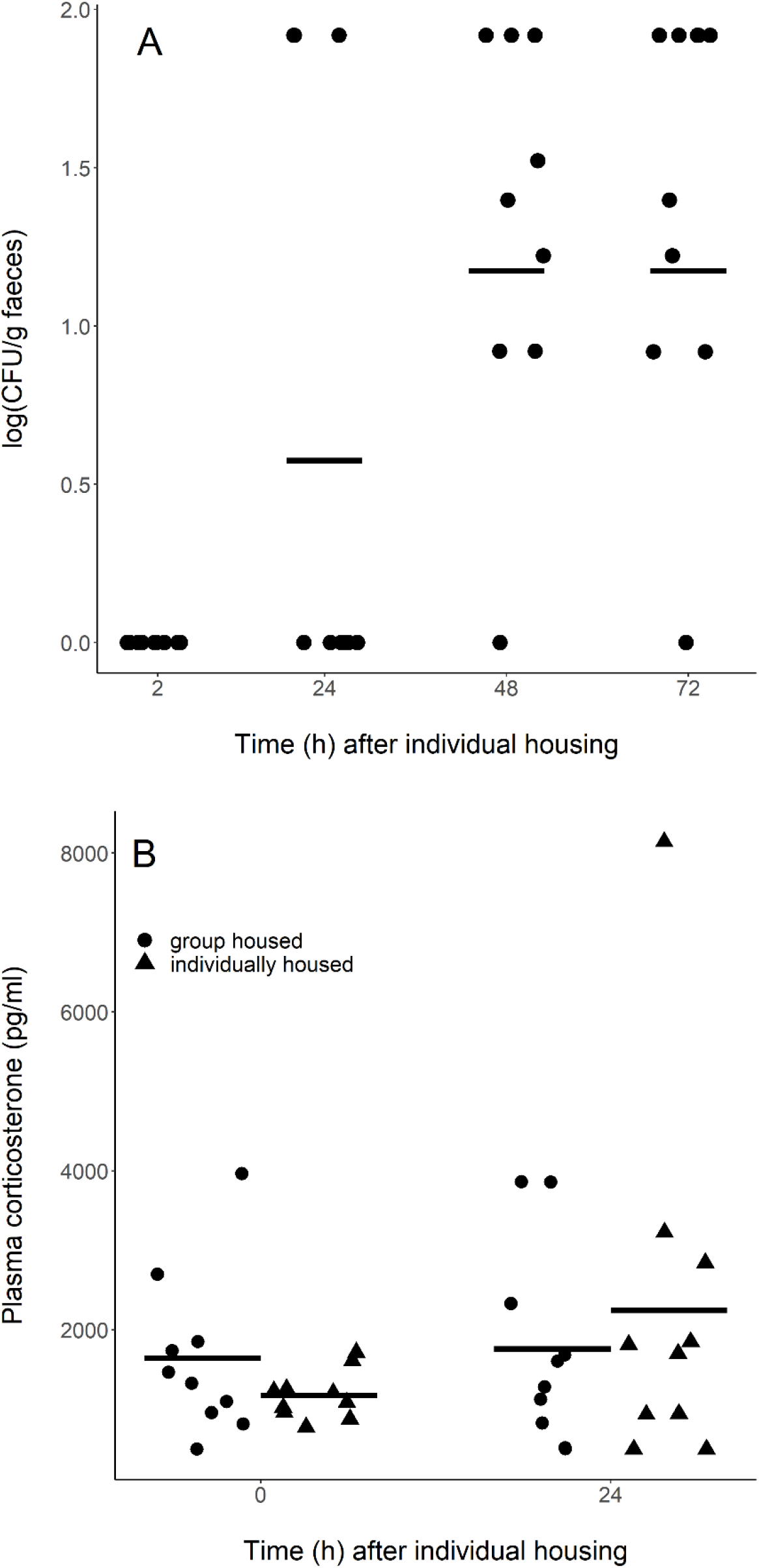
Faecal *Salmonella* shedding (A) and plasma corticosterone levels (B) in pigeons, housed individually or in group. Each symbol represents one pigeon and the horizontal bar indicates the mean value.

## Discussion

### Impaired host condition and reproduction offset by population immunity in a context of pathogen endemism

Successful long-term host-pathogen co-existence entails trade-offs and benefits for both host and pathogen. From the perspective of a host-restricted pathogen, the limited availability of potential hosts supposedly is outweighed by a pathogen competitive advantage through adaptation to a specialized host niche [16,17]. Here, we show that endemism of a host restricted pathogen in the host population comes with a cost for host condition and reproductive success but also with a clear long-term benefit for host disease resistance. *Salmonella* affected body weight and thus body condition in infected pigeons. While more severe during early establishment of infection, this effect persisted at least one year after pathogen introduction. The observed negative impact of bacterial pathogen endemism on host reproduction resembles effects of chronic parasite infections, such as *Plasmodium sp.* infections in blue tits (*Cyanistes caeruleus*) [18,19], schistosome infections in snails [20] and *Toxoplasma gondii* infections in mice [21]. Impaired reproduction and weight gain could be explained by increased host investment (e.g. acquired immunity) in controlling pathogen burden [18,21] and by the development of lesions (granulomata) associated with *Salmonella* persistence [22] and should be considered a trade-off for both the pathogen (reduction of the number and condition of suitable hosts) and the host (potential negative impact on population persistence). The host investment in immunity confers protection against paratyphoid disease at population level. The presence of high titers of antibodies in nestlings at day of hatching combined with exposure to *Salmonella* via parental crop content and the lack of *Salmonella*-induced mortality in early life suggest efficient protection by maternal immunity shortly after hatching. A marked decrease of serum antibody levels by 14 days of age coincided with elevated mortality in older nestlings, suggesting an immunity gap between passive and active immunity [23–26]. When translating these experimental findings to a real world pathogen context, where *Salmonella* was found to be widely present in populations of feral pigeons, population immunity is likely to offer a distinct advantage. At population level, the protection against clinical disease and population decline at least partly compensates condition and reproductive costs associated with host-pathogen co-existence.

### Exploiting host characteristics for pathogen transmission and persistence

Maximizing pathogen transmission maximizes the probability of pathogen maintenance in a host population [27]. We here demonstrate that *Salmonella* exploits the host specific physiological trait of crop-feeding to promote colonization of novel hosts. Shedding of *Salmonella* in the crop could only be detected in adult birds crop-feeding their offspring. Since the stress associated with the breeding period resulted in a significant *Salmonella* recrudescence (see further), shedding in the crop may be the mere result of the overall increased *Salmonella* burden in the parent pigeon. This crop-feeding route of pathogen transmission is a pseudo-vertical transmission [28], allowing fast and efficient transmission of the pathogen to the parents’ offspring without the risk of infertile eggs or embryo mortality. Indeed, despite close association with the host gonads, we could not find any evidence of true vertical transmission (incorporation of *Salmonella* in the egg, resulting in an infected hatchling). The association of *Salmonella* with the pigeons’ gonads thus contributes to pathogen persistence in the host, but not significantly to pathogen transmission. Persistence in the pigeon gonads may be explained by the specific niche *Salmonella* occupies here. Immunohistochemistry showed *Salmonella* to be mainly localized extracellularly in the ovarian medulla and within the tubuli seminiferi and between sertoli cells in the male testes. These sites may favor *Salmonella* persistence through their relative inaccessibility for the host’s immune system [29,30]. Despite *Salmonella* being considered a chiefly intracellular pathogen, residing in the host macrophages [1,14], extracellular persistence in niches poorly accessible to the immune system provides evidence for multiple pathogen strategies [31].

### Host stress promotes pathogen dispersal

Natural stress periods such as breeding, moult and social interactions (introduction of new birds to the group) were shown to temporarily increase *Salmonella* shedding in the pigeons with no discernible impact on bird health. Stress is proven to be a trigger for re-excretion of a pathogen by carrier and corticosteroids have recently been proven to drive *Salmonella* recrudescence [32–34]. Such temporary and corticosteroid associated recrudescence of *Salmonella* infection in the presence of novel hosts (newborn nestlings or naive newcomers) is likely to contribute to pathogen maintenance within a host population.

### Conclusion

We have demonstrated that *Salmonella* Typhimurium exploits host characteristics associated with reproduction and stress to establish and maintain endemism in the pigeon population. This entails a reproductive cost for the hosts, which is outweighed by the development of immunity that protects the birds against pathogen-induced mortality and results in a stable pathogen reservoir.

## Materials and Methods

### *Salmonella* Typhimurium faecal shedding and seroprevalence in feral pigeons

*Salmonella* Typhimurium prevalence in feral pigeon populations was estimated by screening cloacal swabs and serum from pigeons in four populations ((Brussels (50,84667/4,35472; n = 30), Antwerp (51,22139/4,39722; n = 35), Bruges (51,20944/3,22528; n = 39) and Louvain (50,8775/4,70444; n = 29)) for the presence of *Salmonella* Typhimurium and antibodies against *Salmonella* Typhimurium respectively. A clinical examination was done to assess the pigeon’s health status and samples were taken after euthanasia (performed for pest control). An alert, active pigeon without any signs of disease was scored 0; inactive pigeons showing signs of ill health were scored 1. A body condition score, adapted from Gregory and Robins (1998) and Møller *et al.* [36], ranging from 1 to 5 was assigned, based on the bird’s general appearance, breast muscle size and *crista sternalis/breast* muscle ratio; 1: skinny, muscle atrophy, clearly noticeable *crista sternalis;* 2: underdeveloped breast muscles, *crista sternalis* noticeable; 3: normal muscle size, *crista sternalis* palpable; 4: well fed, firm, big breast muscles, *crista sternalis* palpable but not clearly noticeable; 5: obese, big breast muscles, presence of subcutaneous fat behind the sternum. Bacteriological analysis of cloaca swabs and an ELISA on serum samples were performed as described below. For the feral pigeons, it was necessary to first determine the Salmonella serotypes present in the faeces. Therefore, tests were performed on possible (‘pink’) colonies, which were confirmed to be *Salmonella* based on their biochemical characteristics: glucose fermentation, absence of lactose fermentation, H2S production, lysine decarboxylation positive and absence of urease activity. Subsequently they were serotyped by slide agglutination, targeting the somatic antigens (Pro Lab Diagnostics, Bromborough, UK).

### Experimental animals

Forty clinically healthy one-month-old pigeons were obtained from a captive breeding colony free of *Salmonella*. The animals were negative for the presence of *Salmonella* in their faeces on multiple samplings at 1-week intervals as well as for agglutinating antibodies and *anti-Salmonella* IgY antibodies (ELISA) in their blood (see below). Sex was determined using a polymerase chain reaction targeting the CHD genes as described by Griffiths *et al.* (1998). They were group-housed in an aviary with a 12 h photoperiod and fed a commercial seed-based diet *ad libitum.* All the experiments were carried out with the approval of the ethical committee of the Faculty of Veterinary Medicine, Ghent University (EC2013/137; EC2014/96; EC2015/01).

### Experimental Design

Two pigeon groups (similar in age and sex) were formed from the same parental population. In the first group (n = 20), an endemic *Salmonella* Typhimurium infection was established and followed during 66 weeks. The second group (n = 20) was not infected and served as negative control, providing baseline data for clinical signs and reproductive parameters. Mechanisms and trade-offs of *Salmonella* endemism were examined by 1) determining *Salmonella* infection dynamics with focus on identifying routes of transmission, both horizontal (faecal shedding, crop feeding) and vertical (egg and/or semen contamination), 2) quantifying the effect of a *Salmonella* infection on pigeon health and reproductive parameters 3), estimating the role of population immunity in protection against clinical *Salmonella* disease and 4) estimating the contribution of stress periods as inducers of *Salmonella* infection flare-ups.

### *Salmonella* infection and disease dynamics in a pigeon group

*Salmonella* Typhimurium varietas Copenhagen PT99 DAB69, isolated from a pigeon and proven to be pathogenic to pigeons [14], was grown in Luria Bertani (LB) broth overnight at 37°C while shaking. At the age of three months, the 40 pigeons were divided in two groups of 20 pigeons each: (i) pigeons inoculated with *Salmonella,* and (ii) pigeons sham-inoculated. Each pigeon of the first group was inoculated in the crop with 1 x 10^8^ Colony Forming Units (CFU) of the bacterial suspension in 1 mL of inoculum. Birds from the second group were sham-inoculated with LB broth and served as a negative control group. After inoculation, the pigeons, their eggs and their offspring were followed for 15 months (including two breeding seasons) to quantify infection and disease dynamics. Birds were examined daily for the presence of paratyphoid symptoms (anorexia, polydipsia/polyuria, diarrhea, weight loss...) and faecal shedding of the *Salmonella* strain. *Salmonella* shedding in the crop of crop-feeding adults was assessed daily by collecting crop swabs from both pigeon groups. Faecal consistency was scored daily as described by Pasmans *et al*. (2008): 0, normal faeces; 1, faeces not well formed; 2, watery faeces; 3, severe diarrhea; 4, hematochezia; 5, absence of faecal production combined with anorexia.

The numbers of *Salmonella* CFU per gram of matrix were determined by plating ten-fold serial dilutions on Brilliant Green Agar (BGA) plates (LabM, Lancashire, UK). If negative after direct plating, the samples were pre-enriched overnight in buffered peptone water (Oxoid, Basingstoke, UK) at 37°C and then enriched in tetrathionate brilliant green broth (Merck KGaG, Darmstadt, Germany) at 37°C. Therefore, within the infected group, faeces was collected on a daily base to obtain the average daily number of CFU for the entire group and subsequently the average weekly number of CFU. Cloacal swabs and crop swabs were plated on BGA agar and investigated for the presence of *Salmonella* as described above.

At the end of the experiment, all *Salmonella*-infected pigeons were humanely euthanized with an intravenous injection of sodium pentobarbital (100mg/kg, Natrium Pentobarbital 20%, KELA, Belgium) and necropsied. The organs (intestines excluded) were scored for the presence of lesions by the same person using the following system: 0, no macroscopic lesions; 1, organ enlargement; 2, presence of small granuloma’s of < 3 mm; 3, presence of granuloma’s of 3-6 mm; 4, presence of granuloma’s of 7-9 mm; 5, presence of granuloma‘s of > 9 mm. The intestines were scored using the following system: 0, no macroscopic lesions; 1, serosal congestion; 2, abnormal content; 3, hemorrhagic content; 4, presence of granuloma’s of <3 mm; 5, presence of granuloma’s of >3 mm. Tissues were homogenized and the number of CFU of *Salmonella* per g tissue was determined as described above.

### Dynamics of circulating antibodies in pigeons with endemic *Salmonella* infection

We used serum anti-*Salmonella* antibodies dynamics as proxy for the acquired immunity response against the *Salmonella* strain in the pigeon groups. To detect anti-*Salmonella* antibodies, a specific eELISA was developed. This ELISA was home-made according to Leyman *et al.* (2011) with some minor modifications. ELISA plates (F96 maxisorp Nunc-immuno plates, Nunc, Roskilde, Denmark) were coated with 140 μL of a suspension containing formalin-inactivated *Salmonella* Typhimurium varietas Copenhagen strain DAB69 bacteria diluted in coating buffer to an optical density (OD) of 660 nm, measured with a spectrophotometer (Ultraspec III®). Serum was diluted 1/1000 and added to the wells (100 μL). Conjugate consisted of a 1/1000 dilution of a goat anti-bird IgY antibody (Alpha Diagnostics International, San Antonio, Texas, USA). All samples were run in triplicate; 2 positive and 2 negative controls were added to each plate. The washing steps in this protocol were carried out using the Wellwash 4 Mk 2 (Labsystems Oy, Helsinki, Finland). The optical density was measured using a Multiskan MS Reader (Labsystems Oy, Helsinki, Finland) with the Ascent Software, version 2.6. Using this ELISA, blood samples (0.5 mL), collected at 2-month intervals were examined for the presence of anti-*Salmonella* antibodies in all pigeons of both the infected and non-infected group during the whole experiment.

### Protection of pigeons from a group with endemic *Salmonella* infection against clinical salmonellosis

To assess to what extent pigeons born in the endemically infected group were protected against a subsequent *Salmonella* infection and to what extent the presence of anti-*Salmonella* antibodies correlated with protection, we randomly removed eight age- and sex-matched pigeons born in the negative control group and eight born in the infected group from their groups and inoculated them with the *Salmonella* strain. The pigeons of between 5-6 months of age were individually housed and a blood sample (0.5 mL) was taken prior inoculation to check for the presence of anti-*Salmonella* IgY antibodies. Each pigeon was inoculated in the crop with 1 x 10^3^ CFU *Salmonella* Typhimurium PT99 DAB69 in 1 mL of inoculum. The numbers of *Salmonella* CFU per gram of faeces were determined and the birds were followed up clinically as described above. At day 14 post inoculation, a blood sample (0.5 mL) was taken and the pigeons were humanely killed. Internal organs were collected and processed for *Salmonella* quantification as described above.

### Reproductive cost of an endemic *Salmonella* infection

Two months after inoculation with *Salmonella*, the dark/light cycle was gradually adjusted to a daily 16 h photoperiod. Both groups of 20 pigeons each were housed in an indoor aviary (5×2.5×2.5m), given water and feed *ad libitum* and were provided with nesting sites and given the opportunity to make nests and start breeding to study the effect of an endemic pathogen on the host’s reproductive success. We collected data in both groups with regard to a set of parameters, related to reproduction, to estimate the impact of an endemic *Salmonella* infection on host reproduction. The time required to proceed to nest building, the time until the first egg was laid and the number of eggs laid was noted for pigeons in both groups. After laying, the eggs were weighed to the nearest 0.01 g and measured (length and width) using a caliper to the nearest 0.1 mm and given a unique number. To assess whether *Salmonella* is incorporated in the pigeons’ eggs (vertical transmission), the first clutch of eggs was examined for the presence of *Salmonella*. Therefore a swab was taken from the egg shell before egg surface decontamination. Subsequently, the egg was opened and the egg yolk and albumen were removed followed by egg shell and membrane homogenization. Each egg matrix was examined for the presence of *Salmonella* using the above described method. To assess the potential role of semen in *Salmonella* transmission between birds, semen was collected 1-2 times a week from the male pigeons in both groups using a lumbo-sacral and cloacal region massage technique [39,40]. Semen samples were examined for the presence of *Salmonella* as described above.

The pigeons were allowed to incubate the second clutch of eggs. The following parameters were noted: total number of eggs per group, clutch size and egg mass. Unfertilized eggs, determined by egg candling every two days, were processed for *Salmonella* culture as described above. At hatching, nestlings were weighed and a blood sample (20 μL) was taken from the *vena jugularis* to determine the presence of maternal IgY using ELISA. The number of unhatched eggs was noted and again these eggs were processed for *Salmonella* culture as described above. From then on, the young nestlings were weighed daily until the age of 28 days (fledging age). Blood was taken at 14 and 28 days of age (0.5 mL) from the *vena ulnaris superficiali*s. Cloacal swabs and crop swabs were taken daily from day 1 until fledging and checked for the presence of *Salmonella*. Additionally, crop swabs were taken from the crop-feeding parents. The offspring was further kept in the parental group. One year later, the F1 generation of pigeons was also allowed to breed and raise young. All procedures were repeated as described above.

### Role of stress during *Salmonella* endemism in pigeons

Periodic stress may be important in maintaining endemic infections through its potential to induce infection flare-ups [41]. Known stress periods in our experiments were the introduction of new animals, the molting period and the breeding period [32,42–44], and we tested whether exposing pigeons to such stressors influenced the *Salmonella* numbers shed in their faeces.

To test for a causal link between stress and *Salmonella* shedding in faeces, we used single housing as a stressor (often used in stress-related research, e.g. see Baker *et al.,* 2019; Dunn *et al.,* 2015) and determined faecal shedding of *Salmonella* and corticosterone levels in the blood. Ten pigeons from the infected group (chosen *ad random*, equal sex ratio) were individually housed. A blood sample (0.5 mL) was taken from each pigeon prior to individual housing and 24 h later, this within 30 sec after capture to avoid any other influence on corticosterone levels. After individual housing, faecal samples from each pigeon were collected (within 2 and after 24 h, and after 48 and 72 h) for bacteriological titration. As a control, blood samples (0.5 mL) were collected at similar time points from pigeons that stayed in their group. Faecal samples were processed as described before to quantify the number of *Salmonella* bacteria.

To evaluate the effect of the introduction of new animals as a stressor on the group, in a second experiment, we examined the fate of naive pigeons upon introduction in the *Salmonella* positive group. For this purpose, 5 randomly selected pigeons (3 males, 2 females), born in the negative control group, were introduced in the infected pigeon group. Cloacal swabs were taken daily and processed as described above. After 1 month, a blood sample was taken and IgY antibodies were assessed using the *Salmonella* specific ELISA as described above.

### Corticosterone analysis

#### Sample pre-treatment and LC-MS/MS analysis

To 50 μL of plasma we added 50 μL of the internal standard (IS) working solution (corticosterone-d8, 10 ng mL^−1^ in methanol) and 400 μL of liquid chromatography-mass spectrometry (LC-MS) grade water, followed by a vortex mixing (15 sec) and equilibration (5 min, room temperature) step. After the addition of 3 mL of diethylether, the samples were extracted for 20 min on a rotary apparatus. The samples were centrifuged for 10 min at 4750 rpm and 4°C. The supernatant was transferred to another tube and evaporated to dryness using a gentle stream of nitrogen (N2; ~ 40°C). The dry residue was reconstituted in 75 μL of LC-MS grade methanol and vortexed for 15 sec, followed by the addition of 75 μL of LC-MS grade water. After vortexing, the sample was passed through a 0.22 μm Millex® Nylon syringe filter and transferred to an autosampler vial. A 10 μL aliquot was injected onto the liquid chromatography-tandem mass spectrometry (LC-MS/MS) instrument. The LC-MS/MS system consisted of an Acquity H-Class Quaternary Solvent Manager and Flow-Through-Needle Sample Manager with temperature controlled tray and column oven in combination with a Xevo TQ-S® MS/MS system, equipped with an electrospray ionization (ESI) probe operating in the positive mode (all from Waters, Zellik, Belgium). More details about the LC-MS/MS method can be found in De Baere *et al.* (2015) (*47).*

Quantification of corticosterone in pigeon plasma was performed using matrix-matched calibration curves. The calibrator samples were prepared by spiking 50 μL aliquots of pigeon plasma with corticosterone levels of 0.0, 0.5, 1.0, 2.0, 5.0, 10.0 and 20 ng mL^−1^. The standard working solutions of corticosterone and the IS were directly applied onto the samples, followed by a vortex mixing step. After 5 min of equilibration, the sample preparation procedure was performed as described above. The basal concentration of corticosterone in the plasma samples used for the preparation of the calibration curve was determined in order to correct the corticosterone concentrations in the study samples. Limits of quantification (LOQ) and detection (LOD) were 0.5 ng mL^−1^ and 0.2 ng mL^−1^ respectively.

### Immunohistochemistry

The gonads (ovary, oviduct and testes) were fixed in 4% phosphate buffered formaldehyde and embedded in paraffin for routine light microscopy. The immunohistochemical staining protocol to specifically stain the *Salmonella* bacteria was done as described by Morrison *et al.* (2012) and Van Parys *et al.* (2010) with some modifications (*48, 49).* In short, sections of 5 μm were cut followed by deparaffinization, hydration and antigen retrieval in citrate buffer (pH 6.0) using a microwave oven. Slides were incubated with a 3% H2O2 in methanol solution (5 min) and 30% goat serum (30 min) to block endogenous peroxidase activity and non-specific reactions, respectively. This was followed by an incubation step with a Polyclonal rabbit anti-*Salmonella* O4 antibody, targeting the O4 somatic antigen (1/1000; Pro Lab Diagnostics, Bromborough, UK) for 30 min. A biotinylated goat anti-rabbit IgG antibody (1/500; DakoCytomation, Glostrup, Denmark) served as secondary antibody. After rinsing, the sections were incubated with a streptavidin-biotin-HRP complex (DakoCytomation) and the brown color was developed with diaminobenzidine tetrahydrochloride (DAB, Dako) and H2O2. Counterstaining was done using hematoxylin before dehydration and coverslip placement. All incubation steps were done using a Dako Autostainer apparatus.

### Statistical analyses

#### *Salmonella* infections are endemic in feral pigeon populations

To quantify the presence of *Salmonella* Typhimurium in the faeces of free-living, urban pigeons, we used the truePrev function of the R library ‘prevalence’ [50]. This method allows deriving a Bayesian estimate of true prevalence from apparent prevalence obtained by testing individual samples, using a sensitivity value of at least 0.90 and a specificity value of at least 0.99 [51]. To test whether *Salmonella* presence was related to pigeon health status or body constitution score, we applied linear mixed models (R library lme4, Bates *et al.,* 2015) (*52*) with city of capture as a random effect. Health status or body constitution score were specified as dependent variable while presence of faecal *Salmonella,* presence of serum anti-*Salmonella* antibodies and pigeon sex and age (and their two-way interactions) were included as fixed effects. Health status was modelled using a binomial model, for health score a Gaussian error distribution was used. We adopted a frequentist approach whereby full models (i.e. models containing all explanatory variables considered here) were reduced in a stepwise manner, by excluding the variable with the highest p-value until only p < 0.05 predictors remained. Statistics and P-values mentioned in the text and tables are from the minimal model (all significant terms included), whereas statistics and P-values of non-significant terms were obtained by fitting each non-significant term separately into the minimal model.

### Endemism noticed in the wild can be experimentally replicated, and is sustained mainly by horizontal transmission despite marked pathogen association with the host reproductive tract

To test whether *Salmonella* Typhimurium becomes endemic when introduced into a colony of naive pigeons, and to verify whether exposure to stress events increases *Salmonella* Typhimurium prevalence (and thus helps maintaining long-term endemism), weekly data were collected on the density of Colony Forming *Salmonella* Typhimurium Units (CFU) present in the colony’s faeces. From an epidemiological perspective, CFU density in a given week will likely be correlated with the previous week, and measurements thus do not represent independent data and should be treated as a time-series. We therefore opted to apply an ARIMA (autoregressive integrated moving average) model to analyze temporal dynamics of faecal *Salmonella* shedding. Models were fitted using the ‘auto.arima’ function of the R library ‘forecast’ [53], using AIC values as criterion for model selection. Designated periodic stress events (i.e. breeding periods, molting periods and introductions of new individuals, see methods) and the total number of pigeons in the colony were included as covariates. Variable coefficients were divided by their standard errors to obtain the z-statistics, which were then used to calculate the P-values reported.

To compare trends in body mass of experimental versus control group pigeons, a linear mixed model was used. Body mass was specified as a dependent variable and time period (every second month), treatment (experimental versus control) and their interaction as fixed effects. Individual pigeon identity was always included as a random effect. Model were run with a Gaussian error distribution. In addition to testing overall trends throughout the experiment, we verified at which time periods significant differences between experimental and control group arose using the glht function of R library ‘multcomp’ [54], to obtain P-values corrected for multiple testing.

### *Salmonella* endemism impairs host reproduction but does not affect adult pigeon health

To test whether the experimental inoculation of pigeons with *Salmonella* Typhimurium resulted in impaired reproduction, differences in reproductive parameters between the experimental and the control group were assessed using linear mixed models. Differences between the experimental and the control group were tested for by specifying any of the aforementioned parameters as dependent variable and time period (if applicable), treatment (experimental versus control) and their interaction as fixed effects. Models were run with either binomial or Gaussian error distribution as appropriate, and variables were log transformed when needed to obtain normality of residuals (i.e. Shapiro-Wilk ≥ 0.95).

### Endemism coincides with population immunity offering protection against clinical disease but not infection

To test whether the experimental inoculation of naïve pigeons born in the negative control group resulted in an increase in clinical symptoms, the occurrence and/or severity of several parameters for disease in the experimental (pigeons born in the infected group) versus the (naïve) control group of pigeons was assessed using linear mixed models. These parameters (i.e. faecal *Salmonella* count, body mass, faecal consistency scores, diverse morphological anomalies, organ lesions, *Salmonella* presence in organs and antibody titers; see above) were collected either daily, weekly or at the end of the experiment after euthanasia. Differences between experimental and control groups were tested for by specifying any of the aforementioned symptoms as dependent variable and time period (daily or weekly), treatment (experimental versus control) and their interaction as fixed effects. For organ lesions at euthanasia, only treatment was included as fixed effect. Individual pigeon identity was always included as a random effect. Models were run with either binomial or Gaussian error distribution as appropriate, and variables were log transformed when needed to obtain normality of residuals.

### Stress periods result in flare-ups of infection

Linear models were used to test whether individual housing (stressor) of endemically infected pigeons resulted in an increase in faecal *Salmonella* shedding accompanied by an increase in plasma corticosterone levels. To test whether housing pigeons individually increased faecal *Salmonella* shedding, amount of faecal *Salmonella* was specified as the dependent variable and time as fixed effect (time after individual housing). Differences in mean plasma corticosterone levels between the control and the experimental group were tested by specifying time period (time after individual housing), treatment (experimental versus control) and their interaction as fixed effects. Models were run with a Gaussian error distribution. F-tests were used to test whether the individual housing stress experiment affected the variance in plasma corticosterone in the control versus the experimental group.

## Supporting information

Statistical Analyses Results

## Acknowledgements

The authors would like to thank C. Puttevils, D. Ameye, S. Loomans and S. De Baere for their excellent technical assistance. We are grateful to the authorities of Brussels, Antwerp, Bruges and Louvain for providing feral pigeons. The authors declare no competing financial interest.

## Supporting Information

**S1-7 Tables**. Detailed statistical results of all analyses performed.

**S1 Fig.** Immunohistochemical staining of the ovary (A) and testes (B) of pigeons after experimental inoculation with 1 x 10^8^ CFU *Salmonella* Typhimurium varitas Copenhagen DAB 69.

**S2 Fig.** Temporal dynamics of serum anti-*Salmonella* antibodies in pigeons, experimentally infected with 10^8^ CFU *Salmonella* Typhimurium varitas Copenhagen DAB69.

**S3 Fig.** Serum *anti-Salmonella* antibodies present in chicks from a *Salmonella* negative and a *Salmonella* positive group at day 0, 14 and 28 of age during both breeding periods.

**S4 Fig.** Serum *anti-Salmonella* antibodies present in pigeons from a *Salmonella* negative and a *Salmonella* positive flock (n = 16), before and 14 days after experimental inoculation with 10^3^ CFU of *Salmonella* Typhimurium varitas Copenhagen DAB69.

