## Supplemental Figures for "Host-pathogen co-existence incurs reproductive costs"

**Supplemental Information**

**Figure S1: Immunohistochemical staining of the ovary (A) and testes (B) of pigeons after experimental inoculation with 1 x 10^8^ CFU *Salmonella* Typhimurium varitas Copenhagen DAB 69.** The extracellular presence of *Salmonella* within the gonads is visualized by targeting the O4 somatic antigen resulting in brown colored bacteria.

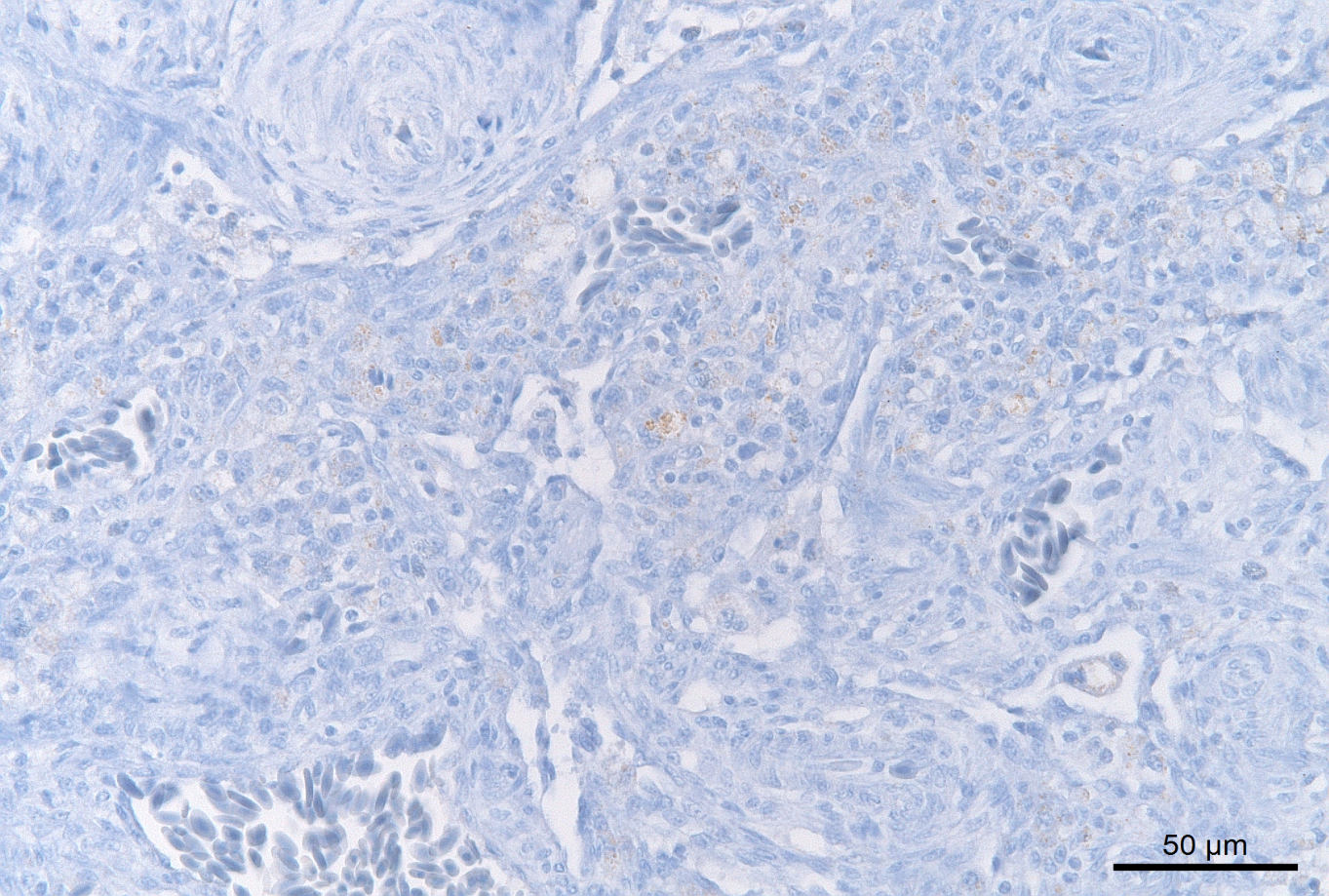

A

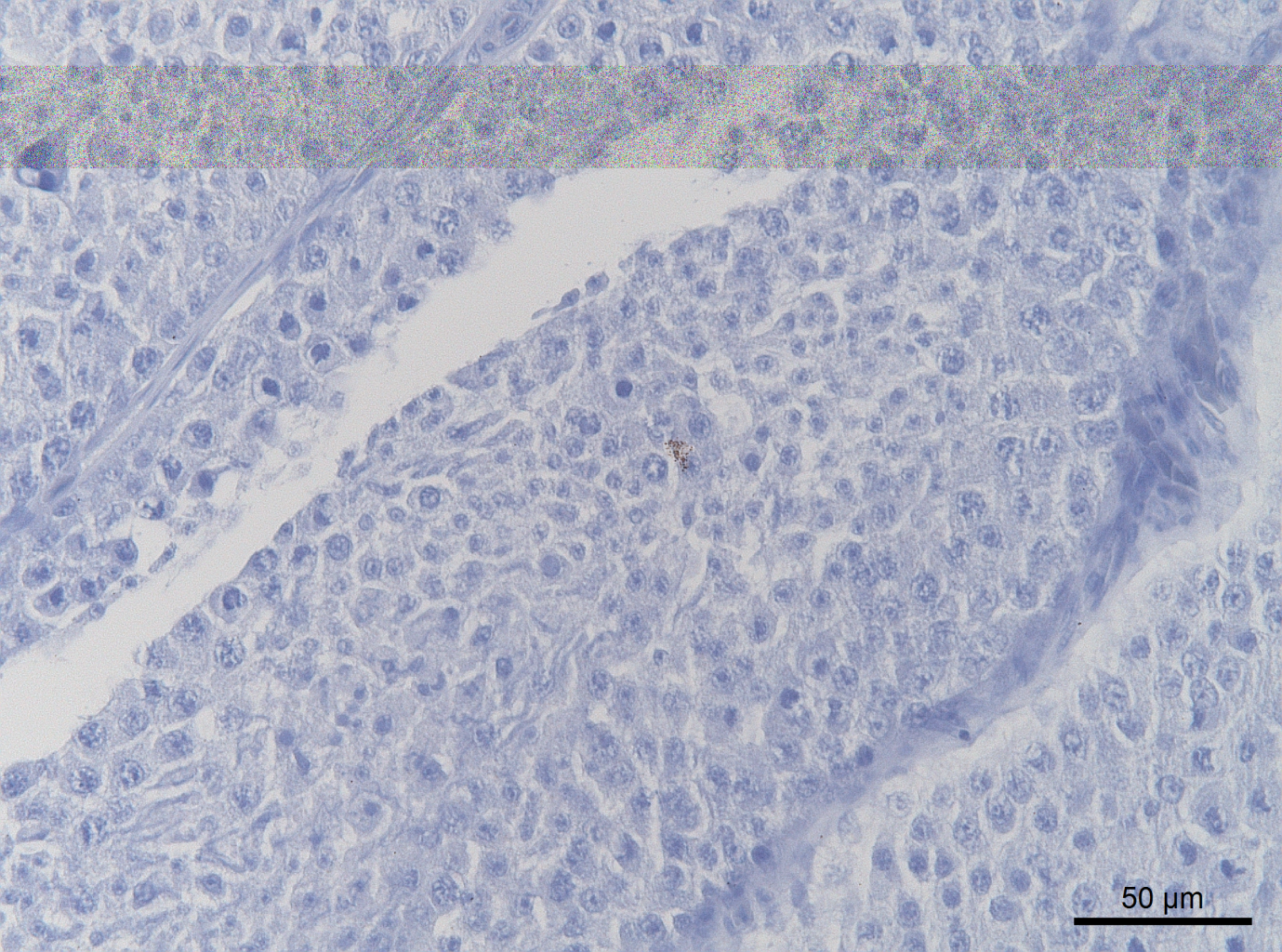

B

**Figure S2: Temporal dynamics of serum anti-*Salmonella* antibodies in pigeons, experimentally infected with 10^8^ CFU *Salmonella*** **Typhimurium varitas Copenhagen DAB69**. Results are presented as mean OD ± SEM. The threshold OD was calculated as 0.2.

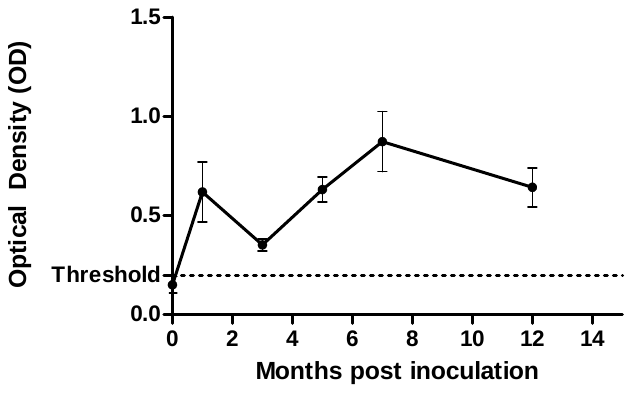

**Figure S3: Serum anti*-Salmonella* antibodies present in chicks from a *Salmonella* negative and a *Salmonella* positive group at day 0, 14 and 28 of age during both breeding periods.** Results are presented as mean ± SD. A significantly higher (p < 0.05) optical density (OD) was noted at hatching (day 0) in chicks born in the experimentally infected group compared to those of the negative control group and is represented by *.

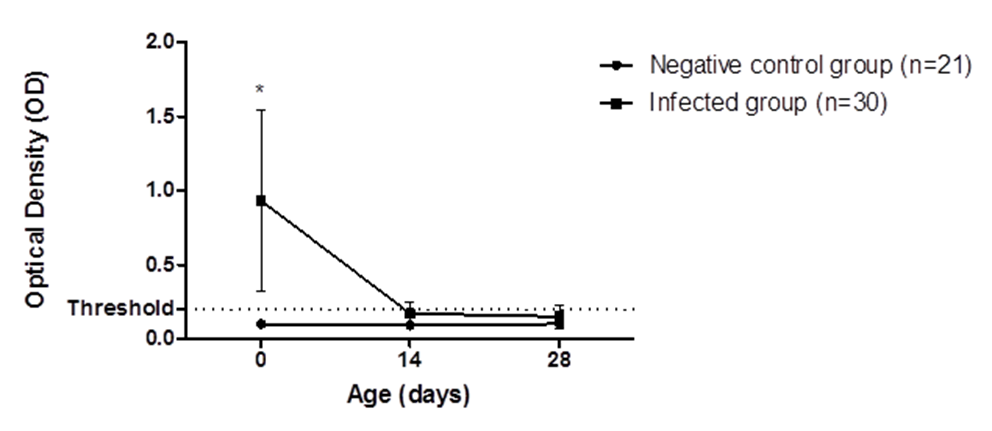

**Figure S4: Serum anti*-Salmonella* antibodies present in pigeons from a *Salmonella* negative and a *Salmonella* positive flock (n = 16), before and 14 days after experimental inoculation with 10³ CFU of *Salmonella* Typhimurium varitas Copenhagen DAB69.** Results are presented as mean ± SEM.
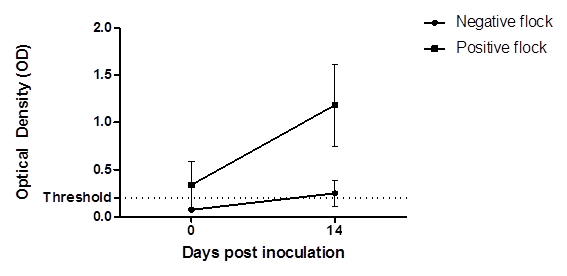
